# Microbial community dynamics during historic drought and flood in the Great Salt Lake

**DOI:** 10.1101/2025.10.23.682790

**Authors:** Alex P. R. Phillips, Amanda E. Lee, Brianne Mortensen, Paulina Martinez-Koury, Sanaz Izadifar, Som Dutta, Bonnie K. Baxter, Amy K. Schmid

## Abstract

Understanding human-driven environmental impacts on microbial community distribution, abundance, and function remains a central challenge in microbial ecology. In particular, the drivers of temporal succession in community membership following perturbation remain unclear. The Great Salt Lake, Utah, bears clear hallmarks of human disturbance, including a rock-filled railroad causeway that sequestered its northern arm from freshwater river influx, leading to localized hypersalination and food web collapse. Following decades of riverine water diversion, the southern arm of this terminal lake reached a historic low elevation during a time of increased climatic change, placing strong environmental pressure on the robust saline ecosystem. Here we use molecular methods to report microbial community composition during this severe drought year at sites across the lake, including where the north and south arm waters contact. At sites of hypersaline water intrusion, we observe surprising stability in north arm community composition, in contrast with strong perturbation in south arm community structure. We use hydrodynamic modeling to pinpoint physical water flow dynamics as a key driver of these community shifts. At saturated salinity north arm sites away from hyposaline water intrusion, abundance shifts were detected in predatory and parasitic taxa, a discovery that reveals surprising ecological dynamics in saturated hypersaline systems. In sum, this study demonstrates drastic hypersaline microbial community shifts during salinity and extreme weather perturbations.

## INTRODUCTION

The Great Salt Lake (GSL) is a natural laboratory for studying extreme environmental impacts on microbial ecology. The largest saline lake in the western hemisphere, this terminal basin features a broad salinity gradient running from the freshwater inputs from mountain snowmelt in the wetlands, to the south arm that accepts the runoff (9-18% salinity), to its salt-saturated north bay (24-34%). As a terminal lake, GSL receives riverine inputs from the watershed draining into the bottom of the basin, but the outflow of water is only through evaporation [1]. The lake levels thus vacillate, and the concentration of salts in the brine are roughly inversely correlated [2]. A history of water dams and diversions in riverine input for consumptive use led to staggering drops in lake elevation at GSL and other terminal saline lakes in recent years [3]. Climate change impacts that enhance drought, such as increased average temperature, decreased precipitation, and a shift from snow to rain have exacerbated the problem [4, 5]. As a result, GSL reached a historic low elevation in late 2022 during catastrophic drought throughout the USA mountain west. The north arm sequestration is due to anthropogenic impacts; around 1960 an unstable railroad causeway that traversed GSL was modified in a way that made a solid barrier to water flow, leading to salinity saturation and a separate north arm ecosystem [6–8]. In 2016, a new breach was constructed, creating a gap in the causeway where boats could traverse. This allowed contact between water originating from north and south lake arms. At the breach, the density difference between the north and south water creates a sharp halocline. In response to the historically low water levels in 2022, the Utah state government built an adaptive rock berm in the breach in July 2022 [9, 10]. Since then, the berm height has been dynamically adjusted to manage salinity flux, especially during seasonal drops in lake elevation when salinity spikes. Federal and state agencies monitor berm impact on water flow, salinity mixing, water quality, nutrients, and lake management [11]. For example, ecosystem monitoring aims to ensure that microbial photoautotrophs in the south arm can support the highly productive food web centered on brine shrimp (*Artemia franciscana*) and brine flies (*Ephydra gracilis)* that feed ten million migratory birds [12, 13]. There are also economic concerns in salinity management, for example *Artemia* cysts are harvested for the aquaculture industry [14]. Economic value of the north arm lies mostly with mineral companies that extract salts and metals from the water. However, the impact of adjustments in the structure and changing hydrology of the causeway on microbial communities across the GSL remains unknown, motivating the current study.

Numerous prior studies have established that the south arm microbial populations shift dramatically in response to temperature, light availability, water inflow, and invertebrate grazing [15–18]. By contrast, the north arm’s saturated brine contains a low diversity, entirely microbial ecosystem, dominated by extremely halophilic (salt-adapted) archaea and bacteria, particularly *Haloquadratum* and *Salinibacter* spp., respectively [17, 19–23]. Evidence suggests that hypersaline environments worldwide (e.g. salt flats) have highly stable compositions compared to those with moderately salinity [24–28]. This may also be true of GSL, although novel lineages unique to GSL of unknown functional capacity were also identified [19, 20]. However, temporal shifts in north arm communities during extreme environmental change remain unclear, especially where the arms interact with each other. To fully understand today’s lake in context of climate change, here we include compositional analysis using improved amplicon reference databases, additional north arm sites, higher resolution time courses, and broader environmental variation. In addition, the identities of the unknown community members, their functional role in the ecosystem, and community mechanisms for maintaining community stability during environmental perturbation such as mixing at the halocline remain unclear.

To address these challenges, here we asked how the GSL community responds to environmental disturbance. Specifically, we hypothesized that relative to north arm samples, breach samples at the mixing point between north and south arm waters will exhibit rapidly shifting microbial community membership and quantity due to highly dynamic water flux. To test this, we compared amplicon sequences of community composition sampled seasonally during an exceptionally variable environmental year in the upper (south arm) and lower (north arm) water column at the new breach vs environmentally stable control sites in each lake arm (Figures 1 and 2). With the increased resolution afforded by this study, a surprising degree of variation was observed in low-abundance north arm taxa and specific variants of major taxa in the context of previously known stability in dominant genera. During the sampling time, the adaptable rock berm in the breach was raised in the summer to block north arm flow and prevent salinity surge in the south arm. Later during the following spring, the berm eroded from high volume snow runoff. As a result, the water turbulence across the sharp halocline at the breach changed over time. Concomitantly, south arm community composition shifted toward that of the north arm but not vice versa. Hydrodynamical models explain these convergence patterns as a function of asymmetric fluid turbulence caused by changes in berm height.

**Figure 1:**
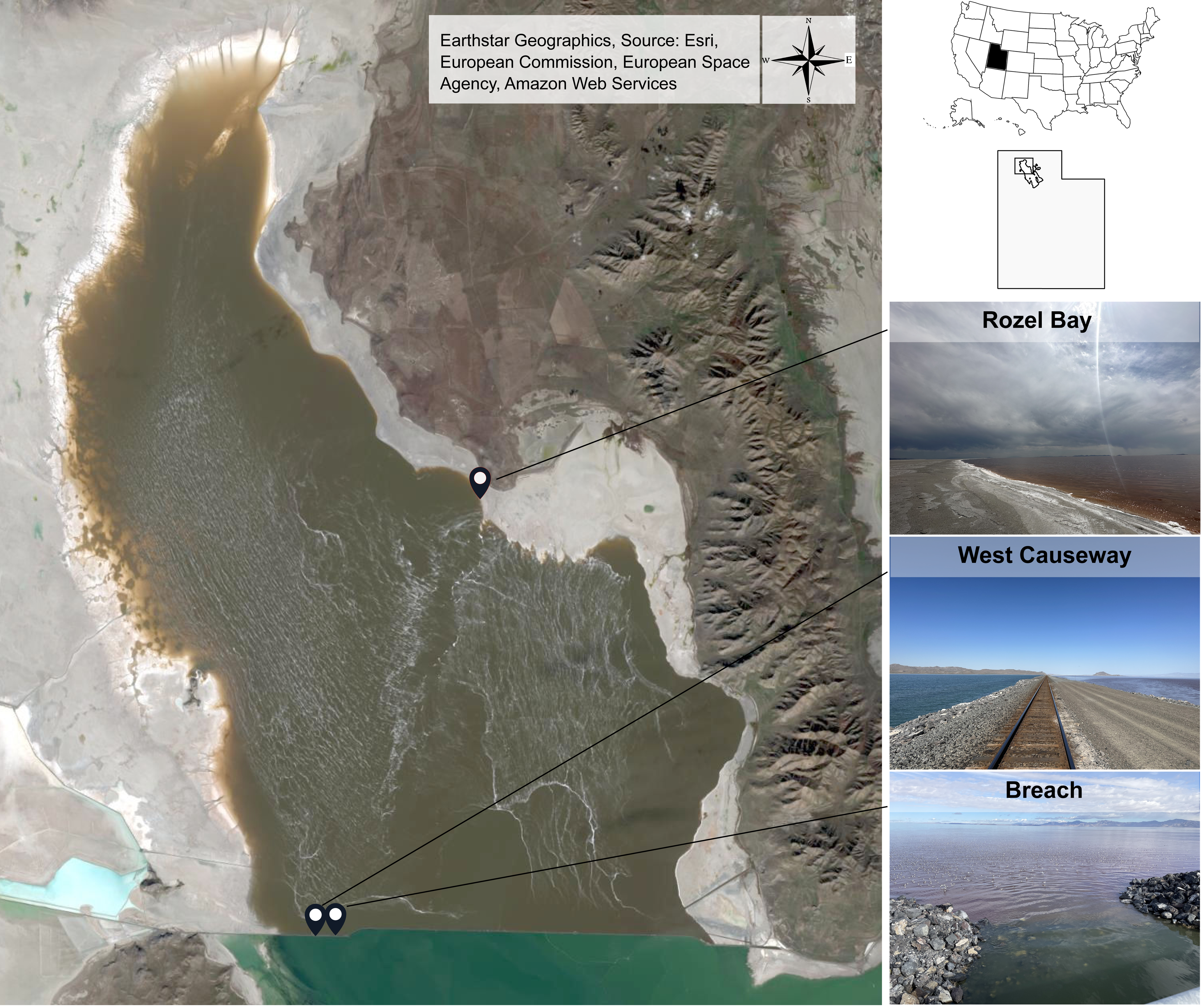
Sampling design. Clockwise from upper left: Satellite image of Great Salt Lake (GSL) with sampling sites indicated by black pins; US map with state of Utah indicated; Utah map with location of GSL shown; picture of Rozel Bay sampling control site; picture of west railroad causeway control site (surface water on north and south sides was sampled); picture of breach, taken from the bridge on the railroad causeway facing north. Compass rose and photo attribution are given in the top right corner of the satellite image. Detailed site information can be found in Table S1.

**Figure 2:**
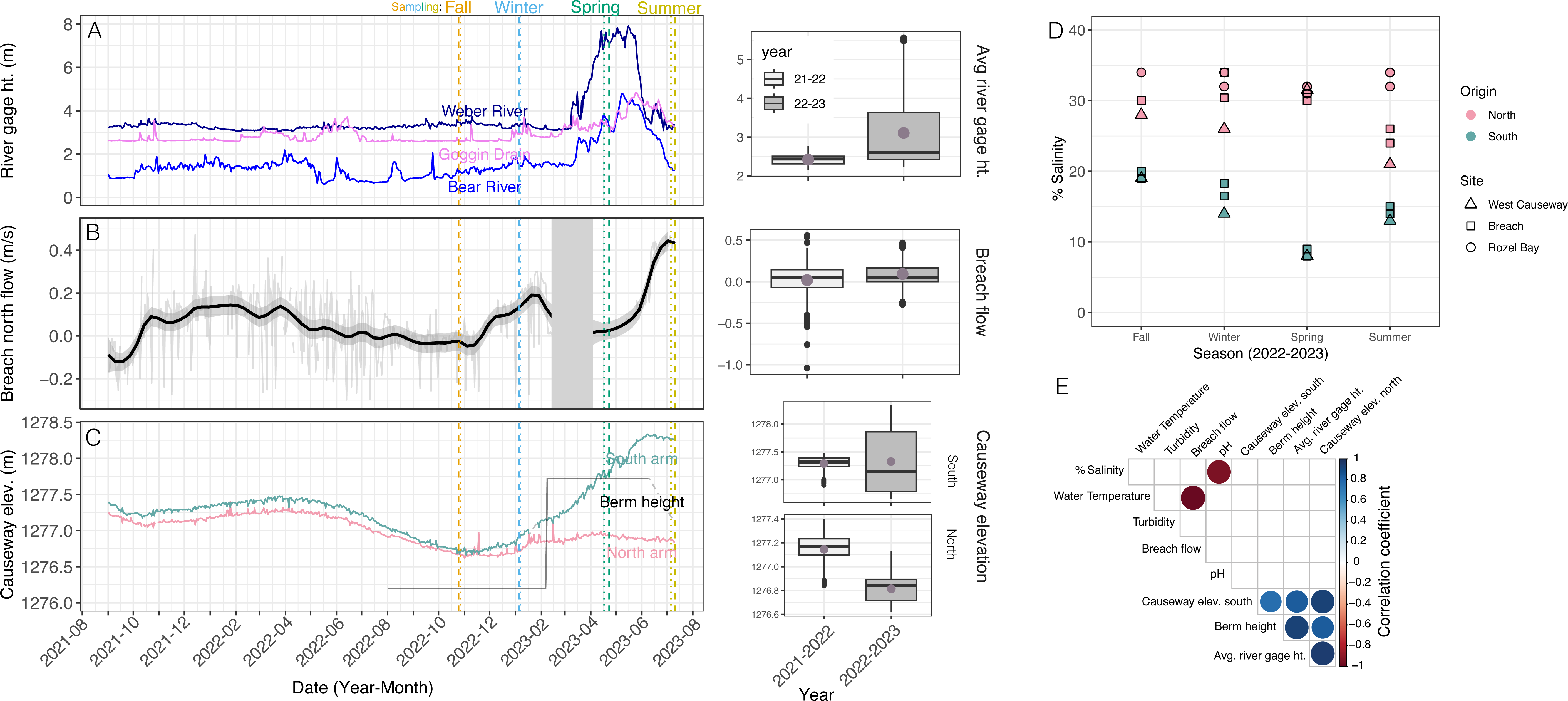
Environmental conditions were variable at the breach relative to the prior year and control sites. Daily values of environmental variables across two years (September 2021 – July 2023) are given in the graphs in the left-hand column. Vertical dotted colored lines represent the time points of seasonal sampling as indicated at the top of the graphs. Vertical dashed lines with long dashes show sampling dates at the causeway, whereas lines with short dashes indicate sampling at the Rozel Bay (1-3 days apart from each other). Summary statistics for each environmental parameter across each year are given in the box-whisker plots at right, where the box length represents the interquartile range (IQR, between first and third quartiles); midline, median; whiskers, 1.5*IQR; black dots are outliers. Lavender points within each box represent the mean of the distribution (mean values are cited in the text). Overbars with asterisks represent a significant difference in the mean of year 1 vs year 2 distribution by Welch’s two-sided unpaired t-test. P-values are cited in the text. (A) Daily river gage heights in meters. Different colors represent data from each of the rivers that flow into the south arm of the GSL, as indicated by text in the corresponding color. (B) Velocity of south to north water flow through the breach in meters per second. Thin grey line represents raw data, heavy black line indicates the loess interpolated and smoothed daily data, and the grey shaded ribbon represents standard error of the loess smoothed mean. Light grey box between February 16, 2023, and April 4, 2023, represents the time period when the sensor was offline and data are interpolated for those time points. (C) Lake water elevation north (pink) and south (teal) of the causeway. (D) Percent salinity at each sampling site. Sites are indicated by different shapes, north vs south arm by different colors indicated in the legend at right. (E) Correlation matrix between abiotic environmental factors across sites and seasons. Significant correlations (*p* < 0.05) between conditions are indicated with a circle. The size of the circle corresponds to the absolute value of the Spearman correlation coefficient, and the color of the circle indicates the Spearman correlation coefficient itself according to the scale bar at right.

## RESULTS

### Historic environmental variation was observed at the breach during seasonal sampling of microbial communities at the Great Salt Lake

To study north arm microbial community compositional change in response to perturbation, GSL water at the breach experimental site was sampled above and below the halocline in each season (Figure 1, GPS coordinates in Table S1, see Methods for sampling details). As controls, samples were taken from surface water at two sites with historical environmental stability [20]: (a) ∼1 km west of the breach from both the north and south sides of the causeway; (b) Rozel Point, ∼25 km north-by-northeast of the breach [29].

During the sampling year, the environmental conditions at the lake transitioned from drought to flood (October 2022-July 2023, Figure 2). Specifically, the GSL watershed experienced the end of a historic drought period in the fall of 2022. In winter 2023, snowpack was double that of 2022 (Figure S1A). GSL water elevation levels surged in spring 2023 as the snowpack melted into the lake basin. During this time, environmental conditions at the breach during the sampling year (September 2022-July 2023) differed quantitatively from the prior year, which was more typical of the breach conditions since its opening in 2016 (see Methods, Figure 2). For example, river inflow gage heights were significantly higher and more variable in the year of sampling than those of the year prior (µ year 1 = 2.42 m, sample year = 3.12 m, *p* < 3.67 x 10^-29^, Figure 2A). Water flow velocity through the breach reflected these river gage trends. In the year prior to sampling, northward water flow velocity through the breach was nearly undetectable but then increased slightly during the winter (range −0.2 – 0.2 m/s, Figure 2B). However, in the sampling year, water flow increased to 0.4 m/s after lake management agencies increased the berm height (Figure 2B &C). Therefore, the overall mean flow velocity across the sampling year (µ = 0.092 m/s) was significantly higher than that in the prior year (µ = 0.019 m/s; *p* < 1.65 x 10^-7^). GSL water elevation at the causeway reflected the trends in flow rate through the breach: both north and south arms reached their lowest recorded elevation at the onset of the sampling year in November 2022 (1276.65 m, Figure 2C). While elevation in the north arm significantly decreased during the sampling year relative to the year prior (from µ = 1277.1 m to 1276.8, p < 2.33 x10^-180^), average south arm water elevation level increased slightly but not significantly between years (µ= 1277.29 m to 1277.32, *p* < 0.310, Figure 2C, right). Although the mean elevation was similar, the range of elevation observed in the sampling year was larger than in the year prior (1276.75 m – 1278.25m). Therefore, water flow at the GSL breach changed significantly during the sample year relative to historical trends. This water flow instability at the breach was reflected in salinity changes. While the north arm control site salinity (Rozel Bay) remained stable near saturation ∼30% throughout the sampling year, south arm salinity reached a record high in the fall of 2022 (∼19%), followed by a gradual decrease on both north and south sides of the causeway as berm height was modulated (Figure 2D). Other environmental parameters such as temperature and pH oscillated seasonally as expected across all sites (Figure S1B and C). As expected, berm height, river gage height, and elevation were strongly and significantly correlated with one another, and pH and salinity were anticorrelated (Figure 2E). However, water flow velocity was anticorrelated with temperature but not significantly related to other environmental parameters, consistent with this parameter being unique to the breach (Figure 2E). In summary, water flow and salinity were variable at the railroad causeway breach relative to control sites. At all sites, conditions varied more during sampling than the typical seasonal vacillation experienced by GSL. These changes are attributable to a complex combination of extreme weather changes in general, and lake environmental management actions at the breach specifically.

### Microbial community diversity dropped significantly during a strong environmental disturbance at the breach

A historic low elevation year followed by a surge of snowmelt were associated with the significant change in environmental parameters observed above (Figure 2). We hypothesized that this alongside human intervention to the breach flow would impact microbial community composition. To test this, we surveyed taxonomic composition by 16S rRNA amplicon sequencing across sites and seasons according to the experimental design described in Figure 1. This yielded 1,084 total distinct amplified sequence variant (ASV) amplicons (Table S1).

Although Shannon diversity of north arm samples was low (∼1.4 to 2.6) and did not vary significantly by site (Kruskal-Wallis rank sum test *p* < 0.3142), significant diversity by season was detected (*p* < 0.0123). In contrast, south arm Shannon diversity ranged from ∼2.2 to 3.5, varying significantly by site (*p* < 0.0015) but not by season (*p* < 0.5425, Figure 3A).

**Figure 3:**
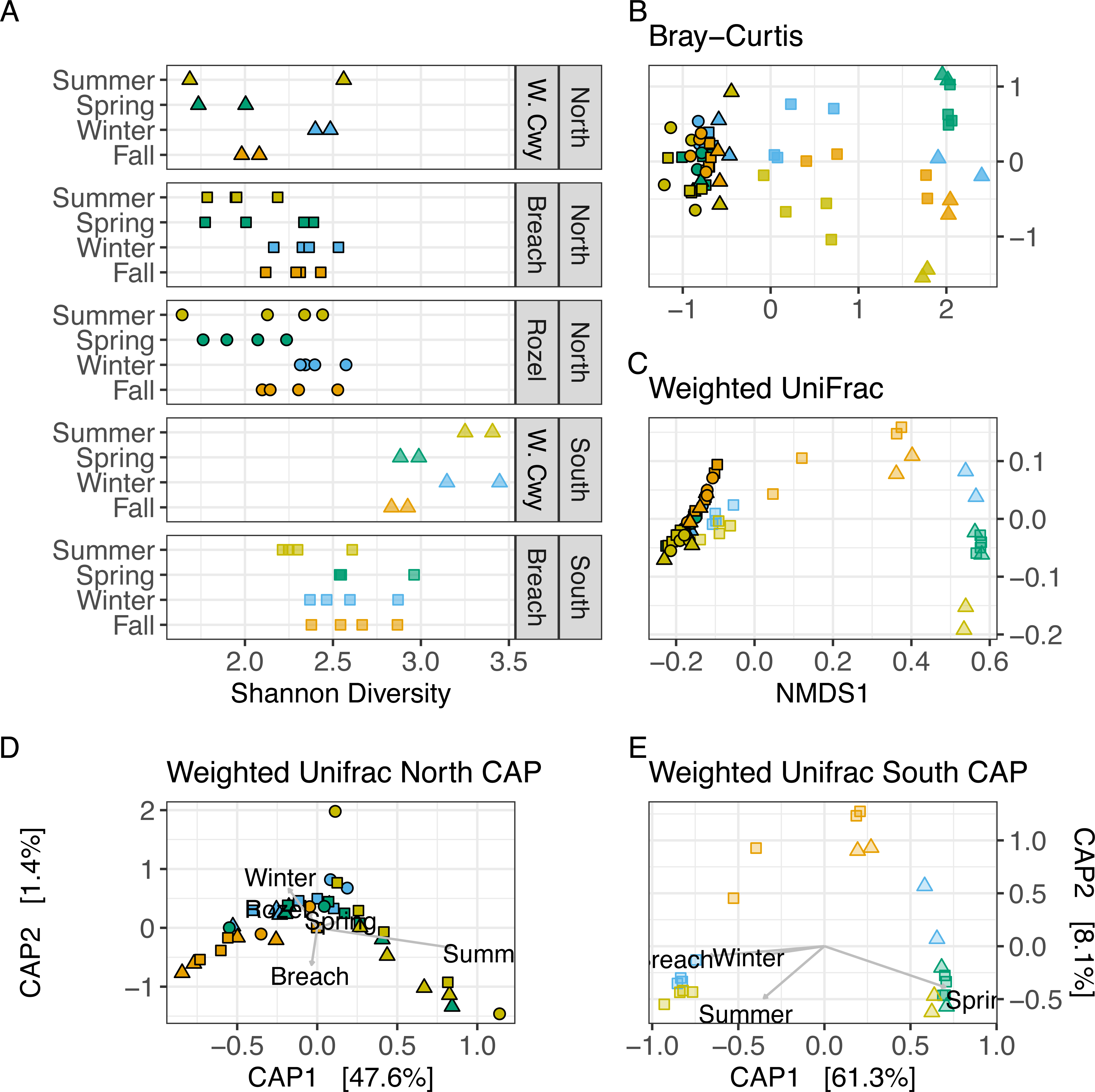
Sample diversity metrics indicate that the south arm breach site community composition loses diversity in every season except spring. (A) Shannon diversity of all samples based on unique 16S amplicon sequence variants, separated by season, site, and lake arm origin. Each point on the graph corresponds to the microbial community at a particular site and season. Points are colored by season and shapes represent sites as indicated in the pictorial legend at right. Duplicate points in each season / site indicate technical replicates. These colors and shapes are consistent throughout all plots in the figure. (B) Nonmetric dimensional scaling (NMDS) plots of sample dissimilarity based on Bray-Curtis distance matrix calculations. (C) NMDS plot of UniFrac distance. (D) NMDS plot of Weighted UniFrac. (D) Constrained analysis of principal components (CAP) plot mapping site and season onto weighted UniFrac dissimilarity plots across north arm samples. (E) CAP plot for south arm samples.

To further investigate whether this change in alpha diversity was due to the drop in diversity at the breach relative to other sites (Figure 3A, bottom panels) beta diversity analysis was performed using the distance metrics Bray-Curtis and Weighted UniFrac. Bray-Curtis distance is proportional to the number of unique shared sequences weighted by their abundances. Weighted UniFrac calculates distance based on both abundance and relatedness of taxa [30]. These metrics showed similar results to alpha diversity (i.e. south arm samples were more diverse than those from the north), but with greater separation in site diversity. In addition, these metrics suggest smaller compositional dissimilarity between communities across north arm sites across seasons compared to those of the south (Figure 3B and C, Table S2). Indeed, water origin had the largest significant effect on community composition according to PERMANOVA analysis (Bray-Curtis F = 63, Weighted UniFrac (WU) F = 78, p < 0.001 for both metrics. P-values and F-statistics for all subsequently discussed comparisons are given in Table S2). Dissimilarity analysis within north arm samples suggested that season is a significant predictor of community composition, with a larger effect size than site (Table S2). In contrast, dissimilarity analysis within south arm samples showed that site had a larger effect on composition than season, although both independent variables had significant effects (Figure 3B-D, Table S2). Specifically, across metrics, contraction in dissimilarity in the south arm Breach site was observed, again suggesting loss of diversity of the community. Breach samples contained sufficient shared taxa with north arm samples to form a separate, intermediate cluster in non-metric dimensional scaling plots (NMDS, Figure 3B-C, Table S2). Site had a larger effect on Weighted UniFrac distance than Bray-Curtis, suggesting that the Breach south arm community shifts significantly toward resemblance of the north arm community in taxonomic composition, relatedness, and abundance (Figure 3C). Surprisingly, in contrast to other seasons, the south arm community remained clustered in the spring across all sites according to Bray-Curtis and Weighted UniFrac metrics (Figure 3B-E). When constrained by season and site, principal component ordination plots showed that the majority (∼60%) of the variation in dissimilarity in the community is explained by the Breach site in fall, winter, and summer vs the spring (Figure 3D and E). Interestingly, the spring season coincided with a substantial increase in the height of the breach dam and a sudden drop in water flow velocity, again suggesting that water flow through the breach could be a key contributor to community composition at this site (see also Figure 2C). Together, these diversity metrics indicate stability across sites with some seasonal differentiation in microbial communities originating from north arm water, even at the highly disturbed Breach site. In contrast, south arm microbial communities lose diversity in breach water during high flow rates, shifting to resemble that of the north arm in every season but spring.

### Microbial community intrusion is asymmetric and dynamic across the breach halocline relative to control sites

To assess specific compositional trends that underlie the observed diversity decrease at the breach, taxonomic analysis of 16S rDNA amplicon sequence variants (ASVs) were compared across sites and seasons for the 1,084 ASVs detected. Across both arms, when the top 200 most abundant ASVs were rank ordered, only 90 ASVs comprised 89.7% of the abundance detected across all samples (Figure S2, Table S3). Further focusing on the top 50 taxa, the top 10 of these explain 87% of the relative abundance regardless of lake arm origin (Figure 4A). These data are in concordance with the low diversity observed by alpha and beta diversity metrics (Figure 3). Only 39 of all detected ASVs were identifiable at the species level, suggesting either poor representation of hypersaline taxa in available databases and/or novel species.

**Figure 4:**
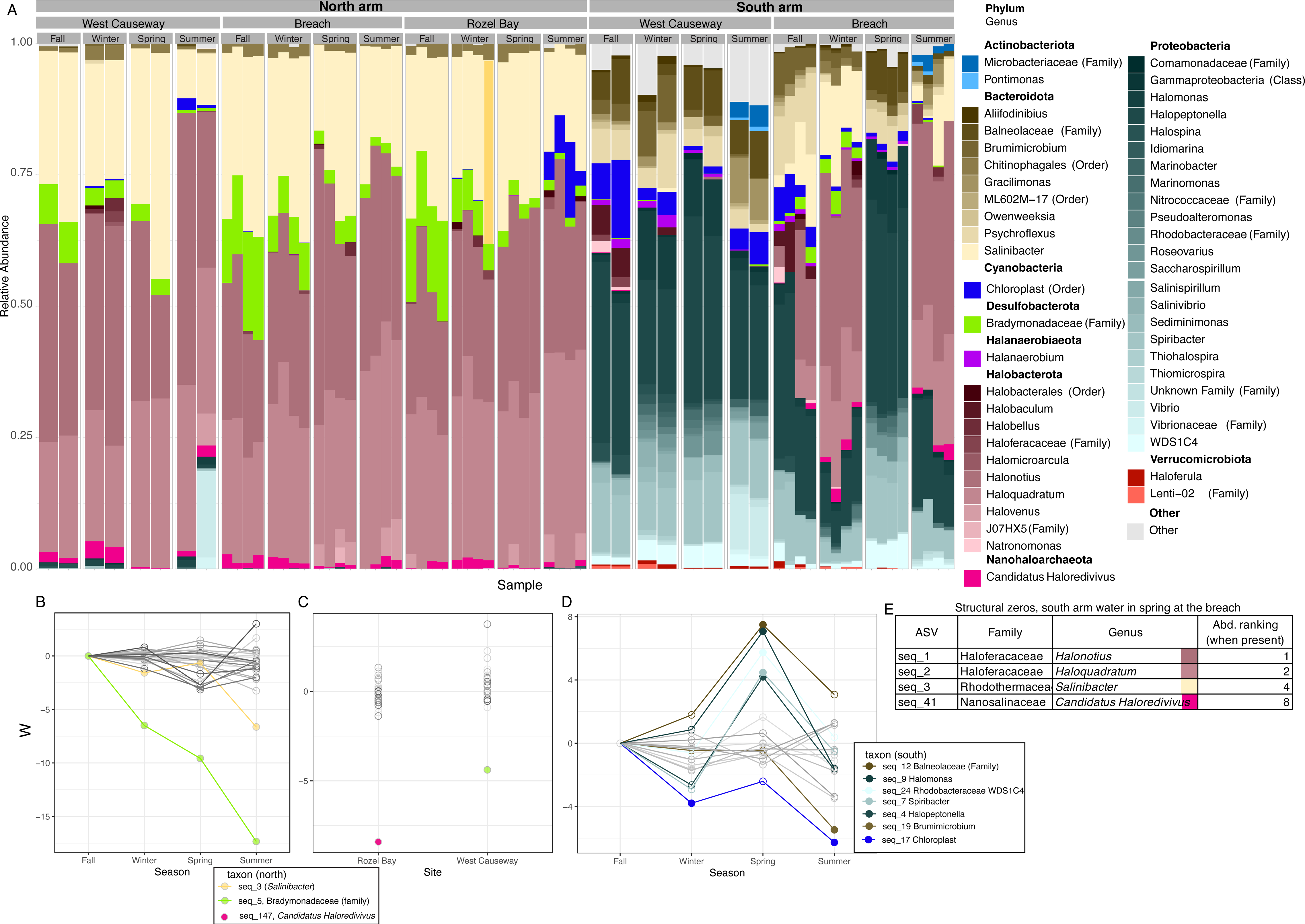
Phylum-level taxonomic composition and relative abundance of the top 200 most abundant taxa across all samples. (A) Sample identities given on the X-axis are labeled in the strips at the top of the bar graph. Biological duplicate samples are paired. Y axis represents relative percent abundance of the top 50 taxa agglomerated by genus within that sample. In the nested bar plot, colors represent phyla, and hues of that same color represent genera as specified in the right-hand color legend. (B) Bias-corrected absolute abundance statistics of specific ASV’s in the north arm. The Y-axis W statistic is set relative to the fall season (see methods for details of the ANCOM-BC model used), representing fold differential abundance of each ASV divided by the standard error in the fall relative to other seasons [32]. Significantly differentially abundant ASV’s are represented by the filled points highlighted in color (see pictorial legend at right for ASV identities). Seasons or sites in which these highlighted ASV’s are not significant are shown by an empty point. All other ASVs that met ANCOM-BC filtering criteria for inclusion in the analysis but not differentially abundant are shown in greyscale. (C) W statistics (log fold change in abundance relative to standard error) for taxa in the north arm breach relative to other sites. (D) W statistics for south arm fall relative to other seasons. (E) Taxa that are abundant in the south arm breach in fall, winter, and summer but not detected in the spring (structural zeros). Right-most column represents the abundance ranking in seasons when the given taxa are detected.

Despite relatively low diversity in the GSL microbial community in general, our analysis further suggests higher stability in dominant taxa in north arm populations than south (Table S1 and S3, Figure 4A). In the north, Archaea (phylum Halobacterota) dominates. Three genera *Halonotius, Haloquadratum,* and *Salinibacter* (domain Bacteria, phylum Bacteroidota) stably dominate the ecosystem across sites and seasons in the north arm, consistent with other hypersaline ecosystems worldwide [31]. Over 15 unique ASVs mapping to each these three dominant north arm genera were detected across seasons, with up to 5 mapping to *Salinibacter* and 4 for *Haloquadratum* (Figure S2). However, only 2 of all 701 ASVs detected in the north arm were differentially abundant according to ANCOM-BC (analysis of composition analysis with bias correction)[32]. Relative to the fall season, ASV seq_3 *(Salinibacter)* showed significant bias-corrected differential abundance only in the summer [adjusted p value (q) < 3.39 x 10^-06^], whereas seq_5 of the Bradymonadaceae family was significantly more abundant in the fall compared to the other seasons (Figure 4B, Supplementary Table S4). Bradymonadaceae are associated with predation in saline ecosystems [33], and they form second-most abundant bacterial group in the north arm after *Salinibacter*. Seq_5 abundance was significantly lower at the west causeway across seasons relative to the breach; therefore, seq_5 differential abundance varied most by season and site relative to all other ASV’s in the north arm (Figure 4C). A specific Nanohaloarchaeal sequence was significantly lower in abundance at the Rozel Bay site than at the breach, although its overall abundance was lower than that of seq_5 (Figure 4A, C). No other north arm ASVs varied by site or season. Together these data suggest that the overall composition of major taxa is stable across seasons in the north arm, with seasonal variation in lower abundance, putatively novel predatory taxa.

In contrast, Bacteria of the phylum Proteobacteria dominate in the south, with microbial diversity and abundance varying substantially by season and site (Figure 4A). The largest fractions of the south arm community were identified as specific variants of *Halopeptonella*, *Spiribacter,* and *Halomonas* bacterial genera. Seven ASV’s demonstrated significant seasonal differential abundance. ASV seq_17, assigned to “chloroplast”, sharply decreased in all seasons relative to the fall (Figure 4D). This ASV showed homology (by nBLAST) to that of trees from the *Pinus* genus, possibly reflecting the accumulation of pine tree pollen in the lake. ASV seq_19 (*Brumimicrobium* genus) decreased significantly in the summer. However, the abundance of the other five ASV’s drastically increased in the spring, especially the three highly abundant proteobacterial genera *Halopeptonella, Spiribacter,* and *Halomonas*. These genera showed stable abundance across other seasons (Figure 4A, D). This significant increase in dominant taxa in spring occurred only at the breach site and coincided with the increase in berm height (Table S4, Figure 2, 4A). The increase in these three proteobacterial ASVs coincided with the complete disappearance of haloarchaeal taxa at the breach in the spring (Figure 4A).

Indeed, ASVs representing the top-most abundant taxa typically found in the north arm water were detected in spring as structural zeros by ANCOM-BC (i.e., they are abundant at the breach in every other season but not detectable in the spring, Table S4, Figure 4E). In contrast, south arm taxa were either undetectable or nearing the detection limit across seasons in north arm breach water (*Halomonas* seq_9 and *Halopeptonella* seq_4, < 0.04% relative abundance, Table S3, Fig 4A). Therefore, consistent with existing knowledge of large-scale seasonal turnover in the south arm water, taxa varied significantly across season. Surprisingly, however, community composition at the breach across the halocline exhibited dynamic asymmetry, with north arm taxa in south arm water but not vice versa. Taken together, these results demonstrate a sharp contrast between the dynamic community composition in the south arm vs overall stability in the north arm across sites and seasons.

### Hydrodynamic flow modeling explains and predicts asymmetric mixing of north and south arm taxa at the breach causeway site

Prior to raising of the berm in August 2022 at the breach, south arm water (∼15% salinity, Figure S2) flowed smoothly northward over the more dense, higher salinity north arm water (∼30% salinity). This density difference stratified the water column, with the layers separated by a sharp halocline (Table S1 tab “metadata”, Figure 5A). Subsequently, the breach berm height was changed at specific points throughout the sampling year: (a) in August 2022, the berm was raised from 1273.5 m to 1276.5 m, resulting in a low flow rate < 0.05 m/s; (b) in February 2023 to 1277 m, reducing northward discharge rate from 0.2 m/s to undetectable levels throughout the spring season; and (c) on June 9^th^, 2023, a section of berm material collapsed, suddenly doubling flow from ∼0.2 m/s to ∼0.4 m/s (Figure 5A) [9]. We reasoned that asymmetry of microbial taxa across the halocline (Figure 4) could result from a physical disturbance of water layers flowing through the breach during changes in berm height.

**Figure 5.**
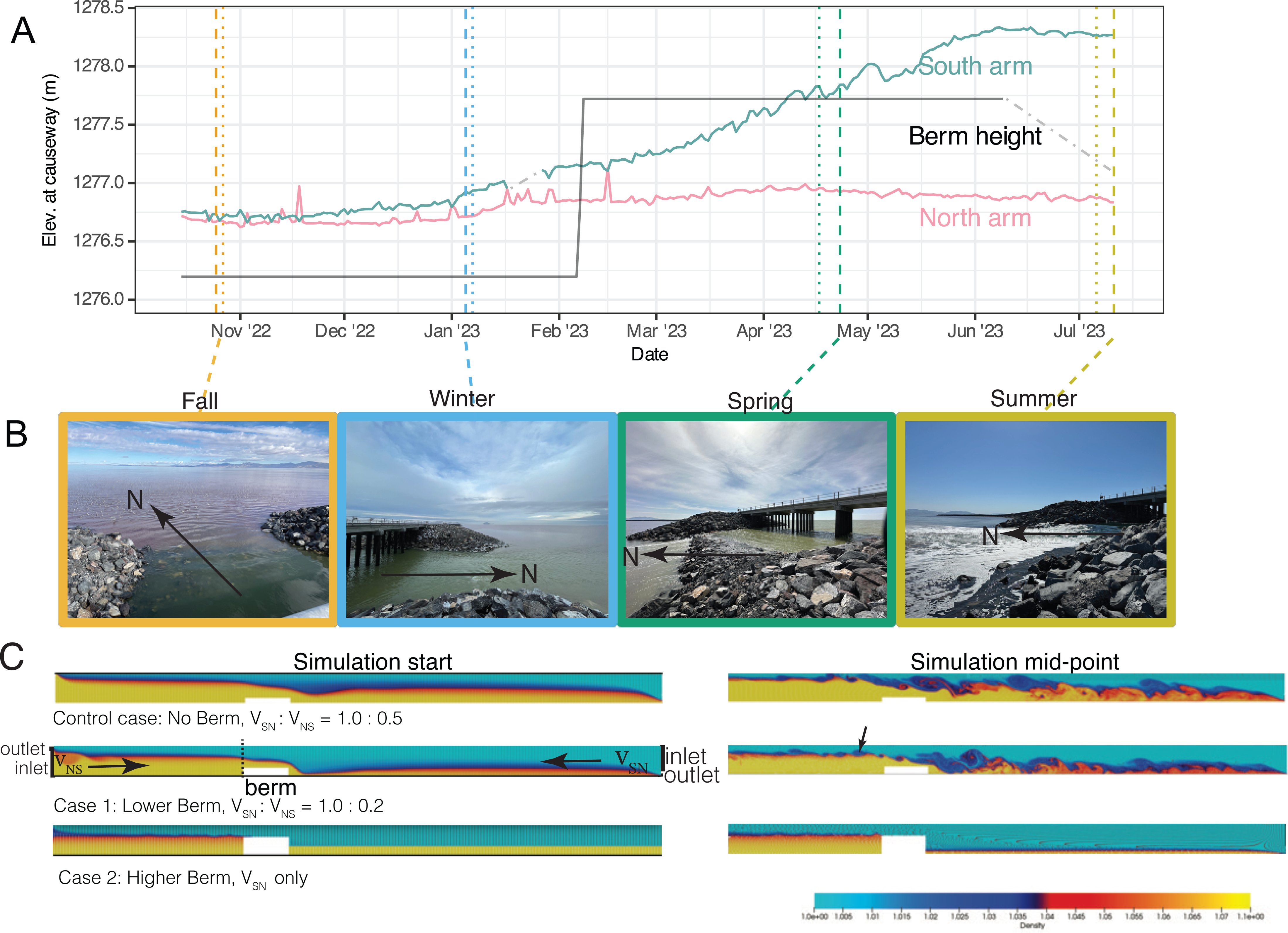
Fluid flow modeling explains the asymmetric infiltration of north arm taxa into the south arm water at the breach. (A) Environmental parameters during the sampling season (same as Figure 2A). (B) Photos of the berm in each sampling season. North is indicated in each image (taken from different angles. Images were captured by A.P.R.P.). (C) Snapshots of model simulation results. Top image represents fluid flow in the case of no berm (lowest-berm) with ratio of south to north flow velocity (V_SN_: V_NS_) as 1:0.5 (control case, not observed during sampling). Middle image represents a low berm (i.e. fall, winter, and summer seasons) with V_SN_: V_NS_ set at 1: 0.2 (see also Methods for modeling details). Lower image represents a berm height above the halocline, V_SN_ only (i.e. spring). Flow direction is indicated by horizontal arrows. Vertical arrow indicates an eddy in south arm flow. Reynolds number (Re, an estimate of turbulence) is set at 10,000 for all the cases. Relative position and scaling of the inlet and outlet points to the simulation are indicated at extreme left and right of the middle snapshot and are conserved across the three cases. Dashed vertical line represents approximate sampling position relative to the berm. Each snapshot was taken mid-way through each simulation. Colors indicate the scale of simulated non-dimensional density (see color legend at bottom right). Movies of simulations for each case are given in Movie S1 (no/lowest berm), S2 (low berm), and S3 (high berm).

To test this, we built a 2-dimensional (2D) high-fidelity eddy-resolved computational fluid dynamic (CFD) model of the flow, based on numerically solving the equations for conservation of mass, momentum and transport of salt with the appropriate boundary conditions (see Methods). We simulated the unsteady dynamics of the buoyancy-driven exchange flow [34] for three scenarios mimicking berm height changes: no berm (hypothetical control case, not sampled), tall berm with low flow (spring), and low berm with high flow (fall, winter, summer). Note that, for the case we are referring to as “no-berm” also has a small, submerged berm. In the no-berm control case, Kelvin-Helmholtz (KH) instabilities [35] generated at the interface of north and south arm layers moving in opposite direction scour the denser north arm water into the less dense south arm layer (Figure 5C, top, Movie S1). This process is further enhanced in the low berm simulation case, where we observed a substantial increase in instabilities introduced into the lower water layer, leading to intensive scouring and mixing with south arm water (Figure 5C, middle, Movie S2). Little to no south arm water was pulled into the denser north arm water. This asymmetric fluid disturbance was further illustrated in the model simulation by the presence of (white) particles carried in the north arm water being entrained and mixed in the south arm water, whereas no (black) particles from the south arm can be found in the north arm water. This happens due to interfacial KH instabilities wrapping the interface into a vortex. As the vortex rolls up, pockets of heavier fluid from the bottom layer are entrained and pulled upward into the lighter fluid. Even though pockets of lighter fluid are occasionally pulled downward into the heavier layer to fill the space, buoyancy opposes this motion, resulting in no mixing of the lighter layer into the heavier layer. However, when the berm was raised higher than the halocline during the early spring of the sampling year, simulated fluid exchange stopped and eliminated mixing (Figure 5C, bottom, Movie S3). Interestingly, a brine layer remained submerged in the south side of the breach in this simulation case, suggesting that mixing at the breach can facilitate microbial dispersal and permanent residence even if the breach were to be sealed at some future point. Taken together, these simulations explain that a low berm coupled with increased flow from riverine input would lead to unidirectional infiltration of water (and therefore hypersaline taxa) resident in the dense hypersaline lower layer into the low-density upper layer.

## DISCUSSION

Here we report a year-long seasonal investigation into the microbial community composition in the hypersaline waters of Great Salt Lake, Utah. We hypothesized that composition and abundance at the interface between the north and south arm waters (breach) would shift during dynamic water flux relative to the hypersaline north arm community. While we largely substantiate this hypothesis, we also find several surprises. Suspected predatory bacteria rise in abundance during cooler seasons in the context of a more stable north arm community (Figure 4). Waters at the breach mix asymmetrically (Figure 5), resulting in a drastic drop in diversity in the low-salinity south arm water at the breach (Figure 3) and providing insight into the negative effects of hypersalination of the south arm. Together, these findings point to possible mechanisms for microbial dispersal and seeding of successive contamination of the south arm with dominant haloarchaeal species.

Therefore, we conclude that microbial community composition in both arms showed varying degrees of dynamic change across the time frame of our study. More specifically, our study recapitulates the north arm community stability observed in previous studies of hypersaline ecosystem composition [19, 20, 24, 25]. However, given the increased temporal resolution of the study design reported here, we also surmised that the north arm community could vary in response to seasonal conditions and south arm connectivity (for example, by lake elevation change and flow through the breach). Consistent with this idea, some specific variants of major genera, and some minor taxa including predatory bacteria, showed shifts in abundance, suggesting niche differentiation, micro-diversity, and the possibility for predator-prey dynamics within these abundant taxa (Figure 2 and 4, Supplementary Tables 2-4). This variation is overlaid with year-round consistency across sampled sites in dominant genera of the north arm surface water microbial community. The co-existence of sequence variants of major taxa in hypersaline systems has been previously demonstrated [36, 37]; however, our study shows for the first time how these vary temporally. Further investigation into the dynamics and metabolic activity of these seasonally varying north arm variants is warranted.

In sharp contrast to the north arm, the south arm breach community was strongly perturbed by changes in water flux. At the breach, asymmetric mixing dynamics across the upper (less saline, lower density south arm origin) and lower (more dense, north arm origin) water layers was observed in every season sampled except spring. Specifically, we observed north arm taxa in water above the halocline (Figure 2E, Figure 4, Figure 5A), but no south arm taxa in denser, high salinity lower-layer water (Figure 5). Using computational fluid dynamics simulations, we found that asymmetric hydrodynamic scouring of the dense north arm water into the surface south arm water indeed explains the asymmetry in the taxonomic distribution detected at the breach (Figure 4, 5B and C). Asymmetric mixing via KH instability has also been observed in the ocean [38]. However, unlike the ocean, a larger difference in density between the two layers of Great Salt Lake water results in stronger asymmetric mixing. We conclude that inherent physical forces across the different water layers significantly perturbed the GSL microbial community. We show that changes in berm height intended by regulatory agencies to control south arm salinity exacerbated water turbulence, thereby further perturbing the resident microbial community (Figure 5).

However, inherent physiological attributes of taxa in the microbial community may also contribute to asymmetric diversity across the halocline. For example, the enhanced range of salinity tolerance of extreme halophiles in the north arm compared to lower salinity south arm-adapted taxa remains a possible and not mutually exclusive explanation for asymmetric taxonomic distribution across the breach halocline. Indeed, extreme halophiles survive at high salinities using an energy-driven sodium/potassium antiporter to maintain a high cytoplasmic potassium concentration (∼4.6 M). To compensate for this salt-in osmotic strategy, they have evolved an acidified proteome [39–41]. Extreme halophiles, especially archaea, therefore dominate in saturated brines and boast a wide salinity tolerance (1.5 – 4.3M NaCl), as we have also observed here (Figure 4) [42, 43]. Haloarchaea may tolerate hypo-osmotic stress in lower-salinity south arm water more readily than salt-out strategists exposed to high-salinity north arm water. For example, the primary producer in these systems, *Dunaliella* spp. algae, maintains osmotic balance through the synthesis of compatible solutes such as glycerol (up to 6-7 M) [44, 45 USA, 46, 47]. Conversely, if salinity levels were to rise in the south arm, haloarchaea would be expected to bloom. Indeed, when salinity neared 20% in the south arm during the fall of 2022, the West Causeway control site exhibited a higher relative abundance of haloarchaea than in any other season at this site (Figure 4A). However, extreme halophiles are not viable below ∼2M NaCl (Sakrikar and Schmid 2021). The drop to 8% salinity in the South-Breach-Spring (Figure 2) coupled with the shut-off of hydrodynamic flow at this time point (Figure 5) could explain the lack of detection of characteristic north arm taxa (Figure 4D). While haloarchaea have been found in south arm water before, previous molecular surveys used separate primers for archaea and bacteria, so changes in their relative abundance have not been systematically measured and directly compared except in this study [18, 19, 48]. It is well established that the moderately halophilic bacteria and eukaryotes of the south arm ecosystem prefer salinities in the range of 8%-15%, and evidence of ecological disruption during the summer and fall of 2022 is mounting [9, 48–51]. Further studies into the salinity tolerance ranges of GSL microbiota under ecologically relevant conditions could help predict, manage, and mitigate future disruption.

Previous studies of brackish water intrusion into estuary-type systems provide precedent for the concept of asymmetry in microbial viability [52]. In these systems, mixed freshwater and saltwater microbial communities converge to a saline-adapted microbial community composition with simultaneous increase in abundance of previously rare taxa that were not adapted to either environment (i.e. niche generalists). However, to our knowledge, saltwater intrusion in ecosystems experiencing the extreme salinity levels examined in this study have not yet been systematically examined. Future research is needed to further test hypotheses regarding mechanisms of microbial survival, interaction, and dispersal in such a dynamic environment. Such investigations would inform prediction of ecological tipping points for microbial successions (e.g. hypersaline archaea population sweep and substantial diversity loss in the south arm ecosystem, Figure 3).

Since the construction of the rock-filled causeway in the 1960s, hypersalination of the GSL north arm resulted in a large-scale ecological succession that was not actively monitored by microbial ecologists with modern methods. Therefore, the current study represents the first, longest running longitudinal survey that used molecular methods to detect hypersaline ecological dynamics in GSL surface waters. We demonstrate diversity loss during hypersalination of the south arm during the 2022 drought. We raise hypotheses to test going forward, including those related to predation, nutrient inflow, and ecological shifts. As climate change-related drought and human consumptive water use continue to threaten sensitive terminal lakes worldwide, the duality of the GSL ecosystem can help to understand the impacts of these perturbations on hypersaline and moderately saline microbial communities concurrently. These data inform the monitoring and management of vulnerable environments and advance our understanding of how ecosystems change in response to factors like desiccation, climate change, and saltwater intrusion.

## METHODS

### Environmental sampling regime and processing

See Table S1 detailed information on sampling sites. At all sites, 1 L of water was sampled from the surface water with 2 x 500 mL sterile plastic Nalgene bottles by hand at control sites, and with a clamp attached to a telescoping pole at the breach. Breach sampling sites were positioned on the northern aspect of the berm with replicate samples taken at the east and west sides of the breach. For north arm water at causeway transect sites, a visual inspection was made to find a location with a layer of north arm water below south arm water. A 500 mL Nalgene polypropylene sample bottle was lowered upside-down through the south arm water layer, into the lower north arm layer, then quickly inverted to sample the dense water, retrieved, and the salinity was verified by refractometer measurements to match that of that of the north arm control sites (∼30%). South arm water was collected from the surface of the upper layer and salinity was verified to match that of the West Causeway control site. At control sites (Rozel Bay and West Causeway), clear surface water (maximum 0.5 m depth) was collected in bottles and salinity measured. At each site, the sampling depth was measured with a ruler. To avoid water cross-contamination at all sample sites, bottles were rinsed once with lake water before sampling, and environmental metadata was collected iteratively and in parallel with sampling. Similar stratification of environmental metadata across the halocline were obtained for this method relative to control sampling with Niskin bottles (Table S5). For all samples, 100 mL lake water was filtered through 0.22 um Sterivex filters (Millipore Sigma, Darmstadt, Germany) with a 50 mL Luer-lock sterile syringe (BD, USA). Filters were flash-frozen in liquid nitrogen, then stored on dry ice during transport (∼2.5 hr). Filters were then stored at −80°C overnight, and DNA was extracted the following day. The causeway and Rozel Bay sites were visited in the morning on separate days no more than five days apart (Figures 2 and 3, methods, Table S2).

### Environmental parameters

A portable refractometer was used to estimate salinity in the field (REED Instruments, Inc., R9600 Salinity Refractometer, 0-28%). Lake water was diluted 1:1 and salinity back-calculated when readings were above 28%. Water clarity was recorded using a turbidity tube with Secchi disk (Eisco Labs, Honeoye Falls, NY). Water temperature was measured with a portable hand-held alcohol or electronic thermometer at the time of collection. GPS coordinates of each site were recorded with the Solocator app (https://solocator.com/) using a mobile phone. U.S. Geological Survey (USGS) data on daily conditions at GSL during the sampling year were accessed via the National Water Dashboard [53] at water monitoring site locations 10010025 and 10010026 for the breach, access date March 25, 2024 (https://waterdata.usgs.gov/monitoring-location/). Additional weather data was sourced from the National Oceanic and Atmospheric Administration retrieved with the Meteostat service (https://meteostat.net/en/). Daily air temperature data was downloaded from weatherunderground.com. Correlation plot shown in Figure 2E was constructed using the corrplot package in R. Detailed weather data used for input to Figure 2 are given at the github code repository associated with this study (https://github.com/amyschmid/GreatSaltLake).

### Nucleic acid extraction

Sterivex filters were opened using a sterilized pipe cutter as in [54]. Filters were transferred to Qiagen PowerWater Pro 5 mL bead-beating kits (Qiagen, USA) and processed according to manufacturer instructions, except that vortexting was run for a total of 10 minutes with cycles of 1 minute on ice and 1 minute vortex bead beating. Lysates were then transferred from the beads by micropipette to fresh 2 mL Eppendorf tubes and processed with the ZymoBIOMICS DNA miniprep kit according to manufacturer instructions (Zymo Research, USA). DNA extracts were stored at −80°C until later use.

### Amplicon sequencing

Community composition in isolated DNA samples was characterized by amplification of the V4 variable region of the 16S rRNA gene by polymerase chain reaction using the forward primer 515 and reverse primer 806 following the Earth Microbiome Project protocol (http://www.earthmicrobiome.org/ and [55–61]). These primers (515F: AATGATACGGCGACCACCGAGATCTACACGCTXXXXXXXXXXXXTATGGTAATTGTGTGYC AGCMGCCGCGGTAA; and 806R: CAAGCAGAAGACGGCATACGAGATAGTCAGCCAGCCGGACTACNVGGGTWTCTAAT) contain barcodes for multiplexed amplification (Table S6). Prior to the selection of these primers, we assessed their coverage of expected organisms by searching their sequence against the Silva database [62, 63] with its TestPrime tool v 1.0 [64] with default settings and verifying matches to hypothesized community members (*Haloquadratum walsbyi* and *Salinibacter ruber*). PCR product concentration was accessed using a Qubit dsDNA HS assay kit (ThermoFisher, Q32854) and a Promega GloMax plate reader. Equimolar 16S rRNA PCR products from all samples were pooled prior to sequencing. Sequencing was performed by the Duke Sequencing and Genomic Technologies shared resource on an Illumina MiSeq instrument configured for 250 base-pair paired-end sequencing runs.

### 16S rRNA amplicon analysis

Sequencing reads were processed with a published 16S analysis pipeline at https://github.com/jianhong/16S_pipeline, based on a framework implemented in Nextflow [65]. Reads were trimmed with Trimmomatic v 0.39 [66] and quality controlled with FastQC v 0.11.9 [67]. Reads were then input to DADA2 [68] v 1.22.0 for read assembly, categorization, and further quality control. Reads were trimmed based on their sequencing quality cutoff, merged, and demultiplexed with default settings. Taxonomy was assigned with the Silva database version 138.1 [62, 63] using Phyloseq [69] v 1.38.0. Once called, the sequences were aligned with MAFFT [70] v 7.475 with the options:--auto –adjustdirection, which chose the strategy FFT-NS-2 and used the progressive alignment method. Approximately maximum likelihood phylogenies were created with IQ-TREE [71] v 1.6.12 by first estimating the model with ModelFinder (TVM+F+R10) and then running with otherwise default options [72]. Taxonomic data visualization and post-hoc analysis was performed with Krona [73] 2.8.1, Phyloseq, fantaxtic [74], and tidyverse [75] packages in using RStudio. The full analysis pipeline snapshot, execution report, and visualization and analysis R code are available in https://github.com/amyschmid/GreatSaltLake. Sequences and raw data can be found in the Sequence Read Archive at BioProject PRJNA1165797.

### Statistics

Correlation analyses were performed in R using the “ordinate” and “distance” functions of the Phyloseq package [69]. PERMANOVA analysis was performed with the “adonis2” function of the vegan package [76]. ANCOM-BC was conducted using the ANCOMBC package in R. ANCOM-BC is specifically designed for detecting the significance and magnitude of abundant taxa in microbiome research [32]. North arm and south arm taxa were analyzed in separate ANCOM-BC models using the ancombc2() function in R. In each analysis, the fall season and breach site were set as the baseline for pairwise comparisons across site + season combinations. An additional, separate analysis of the south arm breach site was performed with the spring season as the baseline in pairwise comparisons to other seasons to detect structural zeros for north arm taxa in south arm water in the breach. For all models, the Holm method was used to adjust p-values for multiple hypothesis testing. W test statistics (log fold change divided by the standard error) for each ASV resulting from ANCOM-BC models are reported in the figures and supplementary material. Specific commands and parameters for all computational and statistical analyses can be found in the github repository associated with the current study https://github.com/amyschmid/GreatSaltLake.

### Fluid flow modeling

Fluid flow through the breach was modeled using the dimensionless form of the Navier-Stokes equations and the advection-diffusion equation, as described previously [34, 77], but extended here to model the buoyancy-driven exchange flow over different berm heights. Simulations were conducted using the high-order spectral element method (SEM) based open-source incompressible Navier-Stokes solver Nek5000 [34, 78]. This high-fidelity solver has been extensively used to successfully model different buoyancy modulated flows in natural [77, 79, 80] and engineered systems [81, 82]. The modeling was conducted in three main steps: (a) writing and derivation of the general equations of mass and momentum conservation, salinity advection-diffusion, and density; (b) abstraction of the problem into non-dimensional space; (c) scaling analysis based on fluid flow from GSL data at the breach to reach the final velocity ratios across the halocline imposed in the simulations [34, 83]. Details of modeling equations, boundary conditions, assumptions, variables, and numerical methods used are given in Supplementary Text File 1.

## ACKNOWLEDGEMENTS

The authors are indebted to the efforts of United States Geological Survey (USGS), especially Hannah McIlwain and Christine Rumsfeld for assistance with on-site sampling of the GSL breach during the early stages of experimental design and implementation. We are grateful for the Utah Department of Natural Resources (DNR), Division of Forestry, and Fire and State Lands for sampling permissions. We thank Great Salt Lake Institute coordinators Georgie Corkery, Jaimi Butler, and Carly Biedul for sampling planning, support, metadata organization, and logistics. We are indebted to many talented Westminster Undergraduates who made technical contributions to GSL sampling and nucleic acid extraction, including Alvin Sihapanya, Anna Jackson, Lauren Rothman, Belen Busquets, and Sierra Watson. This study was funded by National Science Foundation grants 2118274 and 2427099 to AKS; Utah NASA Space Grant NNX15A124H; and a Utah DNR Grant to SD.

## GUIDE TO SUPPLEMENTARY MATERIAL

**Supplementary Text File 1:** Details of hydrodynamic model, including equations and variables.

**Supplementary Figure S1:** Environmental parameters at GSL. (A) Snowfall records from the US Dept. of Agriculture showing snowfall in the year in question. Snowpack peaked at 30 inches / 72 cm snow water equivalent, up from the median peak of ∼16 inches / 40.6 cm. (B) Water temperature at sampling sites during the 2022-2023 sampling year. (C) pH at sampling sites during the sampling year. (D) Correlation plot for relationship between environmental parameters. Dot color corresponds to the extent of correlation according to the color scale bar at right. Dot size represents the absolute value of the correlation coefficient. Blank correlations did not meet the correlation significance threshold of p < 0.05.

**Supplementary Figure S2:** Beta diversity plots with constrained principal components of site, season, and origin, then broken down by arm.

**Supplementary Table S1:** Location (“sampling” subtable) and environmental metadata (“metadata” subtable) across all sites and samples taken for this study. Full 16S amplicon dataset is included with subtables: (a) taxonomoic composition (“taxtab”); (b) sequence abundances by site and season (“seqtab”); (c) raw sequences (“refseqs”).

**Supplementary Table S2:** PERMANOVA results for Origin, site, and season and their effect on beta diversity metrics. Tests were run with vegan::adonis2 in R with 999 permutations.

**Supplementary Table S3:** Top 200 most abundant taxa with relative abundance statistics. Contains data used to make supplementary Table S2.

**Supplementary Table S4:** ANCOM-BC statistical output.

**Supplementary Table S5:** Environmental parameters across the halocline depth profile at the breach using a Niskin bottle (USGS).

**Supplementary Table S6:** Barcode sequences used for demultiplexing 16S amplicon sequences.

## REFERENCES

1. Mohammed IN, Tarboton DG. An examination of the sensitivity of the great salt lake to changes in inputs. Water Resources Research. 2012;48 10.1029/2012WR011908

2. Usgs great salt lake hydro mapper. https://webapps.usgs.gov/gsl/.

3. Wurtsbaugh WA, Miller C, Null SE et al. Decline of the world’s saline lakes. Nature Geoscience. 2017;10:816–21 10.1038/ngeo3052

4. Williams AP, Cook BI, Smerdon JE. Rapid intensification of the emerging southwestern north american megadrought in 2020–2021. Nature Climate Change. 2022;12:232–34 10.1038/s41558-022-01290-z

5. Waddell KaG, W, HDR Engineering, Inc. Final compensatory mitigation and monitoring plan: Union pacific railroad great salt lake causeway culvert closure and bridge construction project. Report. 2016

6. White JS, Null SE, Tarboton DG. How do changes to the railroad causeway in utah’s great salt lake affect water and salt flow? PLOS ONE. 2015;10:e0144111 10.1371/journal.pone.0144111

7. Adams TC. Salt migration to the northwest body of great salt lake, utah. Science. 1964;143:1027–29 10.1126/science.143.3610.1027

8. Madison RJ. Effects of a causeway on the chemistry of the brine in great salt lake, utah. Report. 1970

9. Brown PD, Bosteels T, Marden BT. Salt load transfer and changing salinities across a new causeway breach in great salt lake: Implications for adaptive management. Lakes & Reservoirs: Science, Policy and Management for Sustainable Use. 2023;28:e12421 10.1111/lre.12421

10. U.S.G.S. United states geological survey great salt lake hydro mapper. https://webapps.usgs.gov/gsl/

11. Saccò M, White NE, Harrod C et al. Salt to conserve: A review on the ecology and preservation of hypersaline ecosystems. Biological Reviews. 2021;96:2828– 50 10.1111/brv.12780

12. Baxter BK. The great salt lake food chains: Fragility and resiliency., Salt Lake City, Utah: University of Utah Press, 2024.

13. Sorensen ED, Hoven HM, Neill J Great salt lake shorebirds, their habitats, and food base. In: Baxter BK, Butler JK (eds.). Great salt lake biology: A terminal lake in a time of change, Cham: Springer International Publishing. 263–309

14. Belovsky GE, Stephens D, Perschon C et al. The great salt lake ecosystem (utah, USA): Long term data and a structural equation approach. Ecosphere. 2011;2:art33 10.1890/ES10-00091.1

15. Marden B, Brown P, Bosteels T Great salt lake artemia: Ecosystem functions and services with a global reach. In: Baxter BK, Butler JK (eds.). Great salt lake biology: A terminal lake in a time of change, Cham: Springer International Publishing. 175–237

16. Riddle MR, Baxter BK, Avery BJ. Molecular identification of microorganisms associated with the brine shrimp artemia franciscana. Aquatic Biosystems. 2013;9:7 10.1186/2046-9063-9-7

17. Post FJ. The microbial ecology of the great salt lake. Microbial Ecology. 1977;3:143–65 10.1007/BF02010403

18. Meuser JE, Baxter BK, Spear JR et al. Contrasting patterns of community assembly in the stratified water column of great salt lake, utah. Microbial Ecology. 2013;66:268–80 10.1007/s00248-013-0180-9

19. Tazi L, Breakwell DP, Harker AR et al. Life in extreme environments: Microbial diversity in great salt lake, utah. Extremophiles. 2014;18:525–35 10.1007/s00792-014-0637-x

20. Almeida-Dalmet S, Sikaroodi M, Gillevet P et al. Temporal study of the microbial diversity of the north arm of great salt lake, utah, u.S. Microorganisms. 2015;3:310–26 10.3390/microorganisms3030310

21. Baxter BK. Great salt lake microbiology: A historical perspective. International Microbiology. 2018;21:79–95 10.1007/s10123-018-0008-z

22. Baxter BK, Zalar P The extremophiles of great salt lake: Complex microbiology in a dynamic hypersaline ecosystem. Model ecosystems in extreme environments, Elsevier. 57–99

23. Almeida-Dalmet S, Baxter BK Unexpected complexity at salinity saturation: Microbial diversity of the north arm of great salt lake. In: Baxter BK, Butler JK (eds.). Great salt lake biology: A terminal lake in a time of change, Cham: Springer International Publishing. 119–44. Retreived from 10.1007/978-3-030-40352-2_5

24. Boujelben I, Gomariz M, Martínez-García M et al. Spatial and seasonal prokaryotic community dynamics in ponds of increasing salinity of sfax solar saltern in tunisia. Antonie van Leeuwenhoek. 2012;101:845–57 10.1007/s10482-012-9701-7

25. Di Meglio L, Santos F, Gomariz M et al. Seasonal dynamics of extremely halophilic microbial communities in three argentinian salterns. FEMS Microbiology Ecology. 2016;92:fiw184 10.1093/femsec/fiw184

26. Lee CJD, McMullan PE, O’Kane CJ et al. Nacl-saturated brines are thermodynamically moderate, rather than extreme, microbial habitats. FEMS Microbiology Reviews. 2018;42:672–93 10.1093/femsre/fuy026

27. McGonigle JM, Bernau JA, Bowen BB et al. Robust archaeal and bacterial communities inhabit shallow subsurface sediments of the bonneville salt flats. mSphere. 2019;4 10.1128/mSphere.00378-19

28. McGonigle JM, Bernau JA, Bowen BB et al. Metabolic potential of microbial communities in the hypersaline sediments of the bonneville salt flats. mSystems. 2022;7:e0084622 10.1128/msystems.00846-22

29. Robert Smithson, spiral jetty | exhibitions & projects | exhibitions | dia.

30. Lozupone CA, Hamady M, Kelley ST et al. Quantitative and qualitative beta diversity measures lead to different insights into factors that structure microbial communities. Appl Environ Microbiol. 2007;73:1576–85 10.1128/AEM.01996-06

31. Durán-Viseras A, Andrei A-S, Ghai R et al. New halonotius species provide genomics-based insights into cobalamin synthesis in haloarchaea. Frontiers in Microbiology. 2019;10

32. Lin H, Peddada SD. Analysis of compositions of microbiomes with bias correction. Nature Communications. 2020;11:3514 10.1038/s41467-020-17041-7

33. Mu D-S, Wang S, Liang Q-Y et al. Bradymonabacteria, a novel bacterial predator group with versatile survival strategies in saline environments. Microbiome. 2020;8:126 10.1186/s40168-020-00902-0

34. Rasmussen M, Dutta S, Neilson BT et al. Cfd model of the density-driven bidirectional flows through the west crack breach in the great salt lake causeway. Water. 2021;13:2423 10.3390/w13172423

35. Chandrasekhar S. Hydrodynamic and hydromagnetic stability, New York, NY, USA: Dover Publications, 1981.

36. Ventosa A, de la Haba RR, Sánchez-Porro C et al. Microbial diversity of hypersaline environments: A metagenomic approach. Current Opinion in Microbiology. 2015;25:80–87 10.1016/j.mib.2015.05.002

37. Ventosa A, Fernández AB, León MJ et al. The santa pola saltern as a model for studying the microbiota of hypersaline environments. Extremophiles. 2014;18:811–24 10.1007/s00792-014-0681-6

38. Smyth WD, Moum, J.N. Ocean mixing by kelvin-helmholtz instability. Oceanography. 2012;25:140–49

39. Lanyi JK. Salt-dependent properties of proteins from extremely halophilic bacteria. Bacteriological Reviews. 1974;38:272–90 10.1128/br.38.3.272-290.1974

40. Oren A. Microbial life at high salt concentrations: Phylogenetic and metabolic diversity. Saline Systems. 2008;4:2 10.1186/1746-1448-4-2

41. Christian JH, Waltho JA. Solute concentrations within cells of halophilic and non-halophilic bacteria. Biochim Biophys Acta. 1962;65:506–8 10.1016/0006-3002(62)90453-5

42. Bowers KJ, Wiegel J. Temperature and ph optima of extremely halophilic archaea: A mini-review. Extremophiles. 2011;15:119–28

43. Sakrikar S, Schmid AK. An archaeal histone-like protein regulates gene expression in response to salt stress. Nucleic Acids Res. 2021;49:12732–43

44. Ben-Amotz A, Sussman I, Avron M Glycerol production by dunaliella. In: Mislin H, Bachofen R (eds.). New trends in research and utilization of solar energy through biological systems, Basel: Birkhäuser Basel. 55–58

45. Fendrich C, Schink B. Degradation of glucose, glycerol and acetate by aerobic bacteria in surface water of great salt lake, utah, USA. Systematic and Applied Microbiology. 1988;11:94–96 10.1016/S0723-2020(88)80054-7

46. Elevi Bardavid R, Khristo P, Oren A. Interrelationships between dunaliella and halophilic prokaryotes in saltern crystallizer ponds. Extremophiles. 2008;12:5–14 10.1007/s00792-006-0053-y

47. Oren A. The ecology of dunaliella in high-salt environments. Journal of Biological Research-Thessaloniki. 2014;21:23 10.1186/s40709-014-0023-y

48. Lindsay MR, Anderson C, Fox N et al. Microbialite response to an anthropogenic salinity gradient in great salt lake, utah. Geobiology. 2017;15:131–45 10.1111/gbi.12201

49. Lindsay MR, Johnston RE, Baxter BK et al. Effects of salinity on microbialite-associated production in great salt lake, utah. Ecology. 2019;100:e02611 10.1002/ecy.2611

50. Baxter BK, Zalar P Chapter 4 - the extremophiles of great salt lake: Complex microbiology in a dynamic hypersaline ecosystem. In: Seckbach J, Rampelotto P (eds.). Model ecosystems in extreme environments, Academic Press. 57–99

51. Frantz CM, Gibby C, Nilson R et al. Desiccation of ecosystem-critical microbialites in the shrinking great salt lake, utah (USA). 2023

52. Rocca JD, Simonin M, Bernhardt ES et al. Rare microbial taxa emerge when communities collide: Freshwater and marine microbiome responses to experimental mixing. Ecology. 2020;101:e02956 10.1002/ecy.2956

53. U.S.G.S. Usgs water data for the nation. 2016 10.5066/F7P55KJN

54. Cruaud P, Vigneron A, Fradette M-S et al. Open the sterivextm casing: An easy and effective way to improve DNA extraction yields. Limnology and Oceanography: Methods. 2017;15:1015–20 10.1002/lom3.10221

55. Caporaso JG, Lauber CL, Walters WA et al. Global patterns of 16s rrna diversity at a depth of millions of sequences per sample. Proceedings of the National Academy of Sciences. 2011;108:4516–22 10.1073/pnas.1000080107

56. Quince C, Lanzen A, Davenport RJ et al. Removing noise from pyrosequenced amplicons. BMC Bioinformatics. 2011;12:1–18 10.1186/1471-2105-12-38

57. Caporaso JG, Lauber CL, Walters WA et al. Ultra-high-throughput microbial community analysis on the illumina hiseq and miseq platforms. The ISME Journal. 2012;6:1621–24 10.1038/ismej.2012.8

58. Walters W, Hyde ER, Berg-Lyons D et al. Improved bacterial 16s rrna gene (v4 and v4-5) and fungal internal transcribed spacer marker gene primers for microbial community surveys. mSystems. 2015;1:10.1128/msystems.00009-15 10.1128/msystems.00009-15

59. Apprill A, McNally S, Parsons R et al. Minor revision to v4 region ssu rrna 806r gene primer greatly increases detection of sar11 bacterioplankton. Aquatic Microbial Ecology. 2015;75:129–37 10.3354/ame01753

60. Parada AE, Needham DM, Fuhrman JA. Every base matters: Assessing small subunit rrna primers for marine microbiomes with mock communities, time series and global field samples. Environmental Microbiology. 2016;18:1403–14 10.1111/1462-2920.13023

61. Minich JJ, Humphrey G, Benitez RAS et al. High-throughput miniaturized 16s rrna amplicon library preparation reduces costs while preserving microbiome integrity. mSystems. 2018;3:10.1128/msystems.00166–18 10.1128/msystems.00166-18

62. Quast C, Pruesse E, Yilmaz P et al. The silva ribosomal rna gene database project: Improved data processing and web-based tools. Nucleic Acids Research. 2013;41:D590–D96 10.1093/nar/gks1219

63. Yilmaz P, Parfrey LW, Yarza P et al. The silva and “all-species living tree project (ltp)” taxonomic frameworks. Nucleic Acids Research. 2014;42:D643–D48 10.1093/nar/gkt1209

64. Klindworth A, Pruesse E, Schweer T et al. Evaluation of general 16s ribosomal rna gene pcr primers for classical and next-generation sequencing-based diversity studies. Nucleic Acids Research. 2013;41:e1 10.1093/nar/gks808

65. Di Tommaso P, Chatzou M, Floden EW et al. Nextflow enables reproducible computational workflows. Nature Biotechnology. 2017;35:316–19 10.1038/nbt.3820

66. Bolger AM, Lohse M, Usadel B. Trimmomatic: A flexible trimmer for illumina sequence data. Bioinformatics. 2014;30:2114–20 10.1093/bioinformatics/btu170

67. Andrews S. Babraham bioinformatics - fastqc a quality control tool for high throughput sequence data. https://www.bioinformatics.babraham.ac.uk/projects/fastqc/

68. Callahan BJ, McMurdie PJ, Rosen MJ et al. Dada2: High-resolution sample inference from illumina amplicon data. Nature Methods. 2016;13:581–83 10.1038/nmeth.3869

69. McMurdie PJ, Holmes S. Phyloseq: An r package for reproducible interactive analysis and graphics of microbiome census data. PLOS ONE. 2013;8:e61217 10.1371/journal.pone.0061217

70. Katoh K, Standley DM. Mafft multiple sequence alignment software version 7: Improvements in performance and usability. Molecular Biology and Evolution. 2013;30:772–80 10.1093/molbev/mst010

71. Minh BQ, Schmidt HA, Chernomor O et al. Iq-tree 2: New models and efficient methods for phylogenetic inference in the genomic era. Molecular Biology and Evolution. 2020;37:1530–34 10.1093/molbev/msaa015

72. Kalyaanamoorthy S, Minh BQ, Wong TKF et al. Modelfinder: Fast model selection for accurate phylogenetic estimates. Nature Methods. 2017;14:587–89 10.1038/nmeth.4285

73. Ondov BD, Bergman NH, Phillippy AM. Interactive metagenomic visualization in a web browser. BMC Bioinformatics. 2011;12:385 10.1186/1471-2105-12-385

74. Author. Fantaxtic - nested bar plots for phyloseq data [Computer software]. https://github.com/gmteunisse/Fantaxtic. 2022.

75. Wickham H AM, Bryan J, Chang W, McGowan LD, François R, Grolemund G, Hayes A, Henry L, Hester J, Kuhn M, Pedersen TL, Miller E, Bache SM, Müller K, Ooms J, Robinson D, Seidel DP, Spinu V, Takahashi K, Vaughan D, Wilke C, Woo K, Yutani H. Welcome to the tidyverse. Journal of Open Source Software. 2019;4:1686 doi:10.21105/joss.01686.

76. Author. Vegan: Community ecology package. [Computer software]. https://vegandevs.github.io/vegan/. 2025.

77. Marshall CR, Dorrell, R. M., Dutta, S., Keevil, G. M., Peakall, J., and Tobias, S. M. The effect of schmidt number on gravity current flows: The formation of large-scale three-dimensional structures. Physics of Fluids. 2021;33

78. Fischer PF, Lottes, J.W., Kerkenmeier, S.G. Nek5000 web page. https://nek5000.mcs.anl.gov/

79. Fabregat A, Gisbert F, Vernet A et al. Direct numerical simulation of the turbulent flow generated during a violent expiratory event. Phys Fluids (1994). 2021;33:035122 10.1063/5.0042086

80. Fabregat Tomàs A, Poje, A. C., Özgökmen, T. M., Dewar, W. K. Dynamics of multiphase turbulent plumes with hybrid buoyancy sources in stratified environments. Physics of Fluids. 2016;28

81. Merzari E, Obabko, A., Fischer, P., Halford, N., Walker, J., Siegel, A., and Yu, Y. Large-scale large eddy simulation of nuclear reactor flows: Issues and perspectives. Nuclear Engineering and Design.312:86–98

82. Paul MR, Einarsson, M. I., Fischer, P. F., and Cross, M. C. Extensive chaos in rayleigh-bénard convection. Statistical, Nonlinear, and Soft Matter Physics. 2007;75:045203

83. Dunn D, Crookston, B. M., Phillips, C., Dutta, S., and Neilson, B. Seasonal water and salt cycling in the great salt lake after opening the new causeway breach. Journal of Hydrology: Regional Studies. 2025;59:102332

