## Supplementary figures and images for "Microbial community dynamics during historic drought and flood in the Great Salt Lake"

### Fig S1

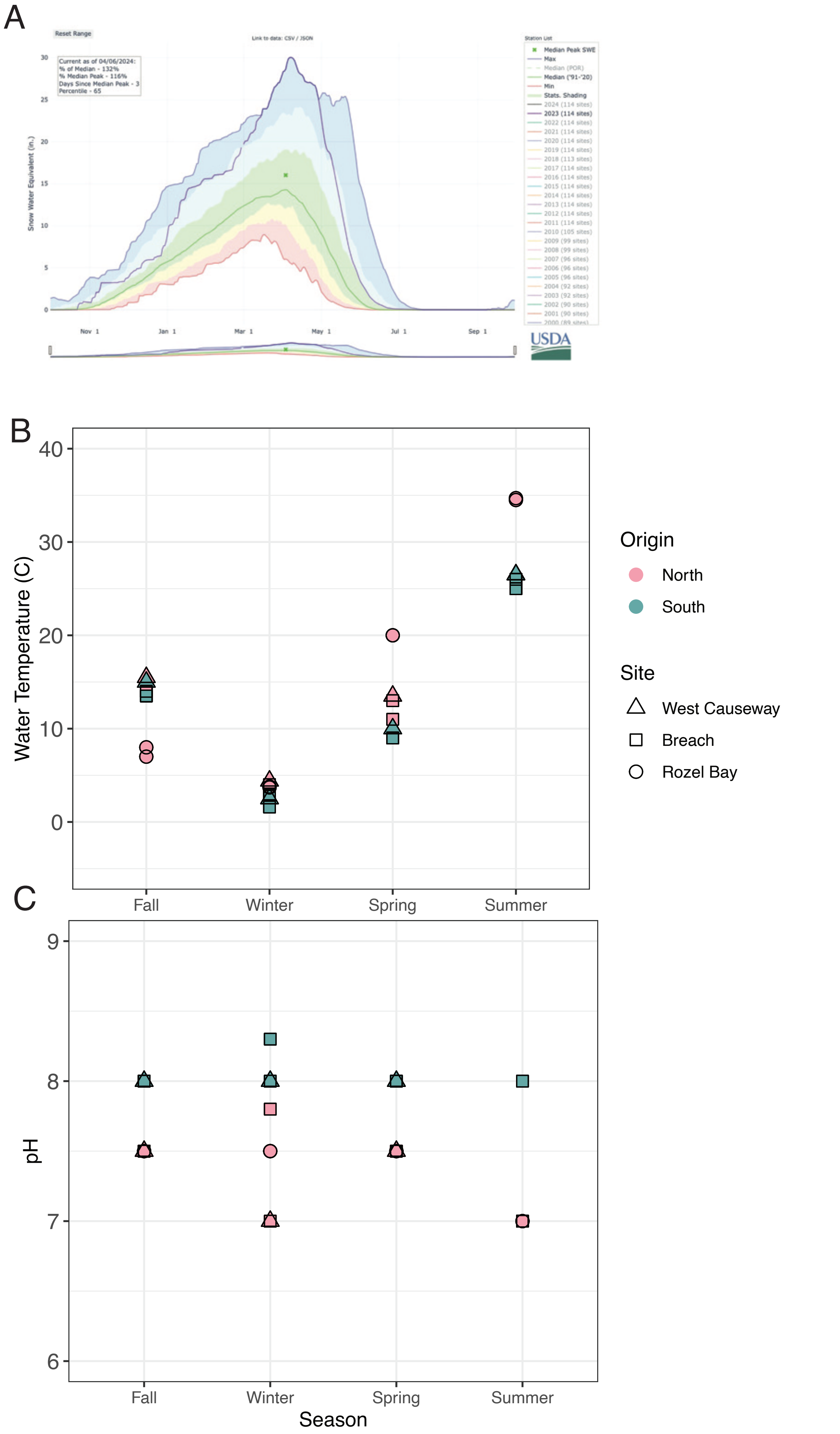

### Fig S2

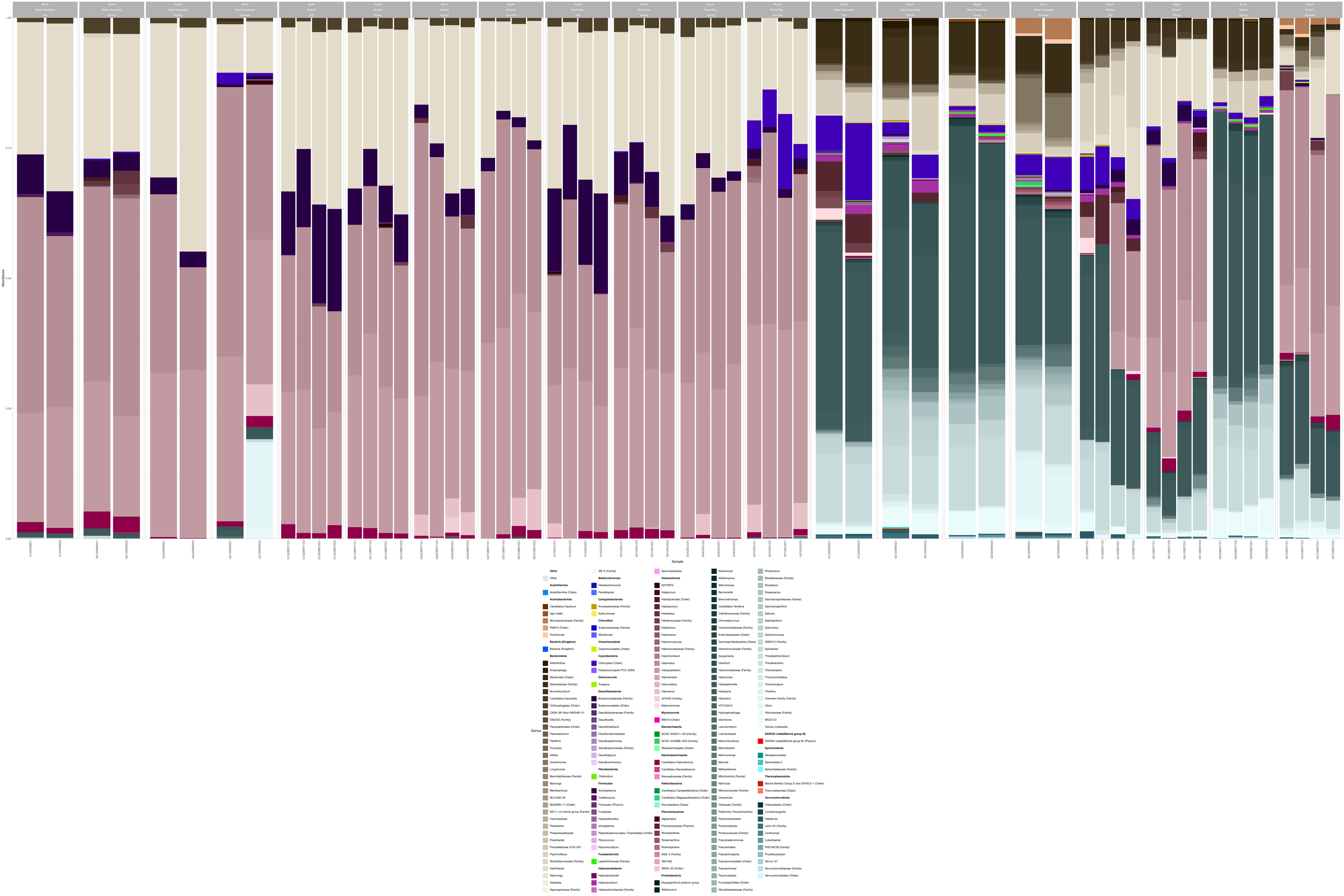
